# Juvenile hormone regulation on the flight capability of *Bactrocera dorsalis* (Diptera: Tephritidae)

**DOI:** 10.1101/867085

**Authors:** Peng Chen, Min Chen, Hui Ye, Ruiling Yuan, Chunhua Du, Su Ping Ong

**Affiliations:** Yunnan Academy of Forestry and Grassland, Kunming 650201, China; Biocontrol Engineering Research Center of Plant Disease & Pest, Yunnan University, Kunming 650091, China; Southwest Forestry University, Kunming 650224, China; Forest Research Institute Malaysia (FRIM), 52109 Kepong, Selangor, Malaysia

**Keywords:** *Bactrocera dorsalis*, Tephritidae, juvenile hormone, flight muscle, chromatography

## Abstract

The oriental fruit fly, *Bactrocera dorsalis* (Hendel) (Diptera: Tephritidae), is considered a major economic threat in many regions worldwide. In order to better understand the flight capacity of *B. dorsalis* and its physiological basis, the functions and regulatory roles of juvenile hormone (JH) in the flight muscle of *B. dorsalis* were studied under a controlled environment. JH titer of *B. dorsalis* varied with age and sex. Females, irrespective of age, have higher JH than males for ovarian development and maturation in addition to better flight capabilities. The flight duration and distance of both males and females increased with the gradual increase of JH titer after adult emergences. JH titer peaked in 15-d-old adult and declined subsequently with age. Flight activity stimulated the production of JH as adults flown for 24 hours on the flight mills have the highest JH titers compare to adults tethered on shorter flight durations. Furthermore, JH III-treated adults were able to perform long-duration and long-distance flights. The mutual reinforcement of JH and flight activity provides fundamental understanding on the physiological aspects of the flight capability and dispersal, which facilitates strategies for the long-term control of this destructive pest.

## Introduction

Juvenile hormone (JH) is a sesquiterpene compound synthesized via mevalonate pathway in the corpora allata (CA) of insects, which regulates insect metamorphosis, growth and development, reproduction, and flight (Tu et al. 2006). The role of JH in regulating insect flight was discovered in the 1970s (Davis 1975). Among the eight JHs that have been identified, JH 0, JH I, JH II, JH III, 4-methyl-JH I, methyl farnesoate, JH III bisepoxide and JH III skipped bisepoxide, JH III is the most common form in various insects (Mauchamp et al. 1999, Steiner et al. 1999, Gilbert et al. 2000; Kotaki et al. 2009).

Different species of insects have different physiological responses to the same hormone while the same species of insect can also exhibit diametrical opposite responses due to the differences in the concentrations of JH (Rankin 1991, Zera 2004). For example, studies on the brown planthopper and wing-dimorphic crickets have shown that the increase of JH concentration led to the degradation of flight muscles, followed by inhibition of flight activities (Tanaka 1994, Dai et al. 2004, Bertuso and Tojo 2002, Zhao and Zhu 2013). In contrast, increase of JH titer in migratory insects such as the convergent lady beetle and western corn rootworm was associated with increase in flight capabilities (Coats et al. 1987, Rankin and Rankin 1980). In addition, synthesis of JH in the CA was reportedly high after the flight activity of the insect (Li 2004, Jiang et al. 2011).

The strong flight capability of *Bactrocera dorsalis* (Hendel), a destructive agricultural pest, coupled with suitable environmental conditions had enabled its long-distance dispersal and rapid northward range expansion in mainland China (Yan 1984, Liu 2005, Chen et al. 2007). Flight behaviour- and performance-related researches have aimed to elucidate its pattern of invasiveness and effective control techniques (Chen et al. 2007, Yuan et al. 2014, Chen et al. 2015, Chen et al. 2017). Nevertheless, many aspects including hormonal regulation on *B. dorsalis* flight mechanisms remained unknown. This study is a novel approach in the quantification of JH in *B. dorsalis* flight muscle to examine its relationship and implications on the flight capability and ovary development in *B. dorsalis*.

## Materials and Methods

### Test insect

*Bactrocera dorsalis* were obtained from colonies reared in the Forest Protection Institute, Yunnan Academy of Forestry and Grassland. The colonies were maintained at 25°C, 60% relative humidity (RH) and a photoperiod of L12:D12.

### JH extraction

JH was extracted using the protocol of Gharib and Reggi (1983) for optimal extraction. The thorax of *B. dorsalis* was cut off, rinsed with ultrapure water (Dura Pro, United States) to remove surface impurities prior to extraction, dried with a Whatman filter paper and weighed on an electronic balance (Sartorius). 0.1 mg of the sample was homogenized in 1 ml of solvent consisting of hexane-methanol mixture at the ratio of 2:1 and centrifuged at 10 000 rpm for 10 min to separate the hexane phase from the methanol phase. The methanol phase was further rinsed with 600 μl of the hexane-methanol (2:1) solvent and the two hexane fractions were pooled. To optimize the extraction of JH, the methanol phase was rinsed again with 500 μl of hexane and centrifuged at 10 000 rpm for 10 min. All the hexane fractions were then pooled and centrifuged at 10 000 rpm for 10 min to remove impurities and insoluble. The supernatant was pipetted and stored at −40°C. Analytical grade methanol and hexane were obtained from Beijing Chemical Reagent Co., Ltd.

### JH quantification

JH titer in the flight muscle of *B. dorsalis* was determined using high performance liquid chromatography (Agilent 1100 HPLC System, Agilent Technologies) with Diode Array Detector (DAD). A reversed-phase column, Diamonsil® C18 (250 mm × 4.6 mm, particle size 5 μm) (Dikma Technologies, Inc.) was selected as the stationary phase. The samples were dried using the nitrogen blow-down method and added with methanol-water mixture (80:20 v/v) as the mobile phase, to a volume of 30 μl. The injection volume was 10 μl at a flow rate of 0.8 ml/min. These combinations were selected to provide the optimum conditions for the separation of JH III. A UV detector at a wavelength of 220 nm was used with a column temperature of 25°C. The retention times and peak areas of the samples were quantified and compared with the JH III reference standard.

For the preparation of a calibration curve, JH III reference standard (Sigma-Aldrich, St. Louis, Missouri) was diluted into a series of concentrations using methanol as a solvent. Five standard solutions of 1, 10, 25, 50 and 100 mg/ml were prepared. HPLC analysis was performed on the standard solutions to obtain a calibration curve to quantify the concentrations of JH III in the test samples.

### JH titers, flight capability and ovarian developmental processes in *B. dorsalis* adults of different ages and sexes

JHs of six male and six female adults of the same age and similar size were assayed at 5, 10, 15, 20 and 25 days old, respectively, according to the procedures described earlier. JH extractions from the flight muscles of the test insects were done between 2 –3 p.m. for each replicate to ensure consistency of the titers. After JH assaying, the developmental stages in the ovary of tested females were determined based on the morphological structures and developmental characteristics as follows: previtellogenic stage (I), vitellogenic deposition stage (II), expectant stage of mature eggs (III), peak stage of oviposition (IV) and last stage of oviposition (V) (see Chen et al. 2014).

Another batch of six male and six female individuals of the same age and similar size were tested for their flight capabilities at 5, 10, 15, 20 and 25 days old, respectively. Each individual was used only once throughout the experiment. Each flight test was conducted for 13 h and the total number of flight mill revolutions including the flight characteristics of the test individuals (e.g. duration, distance, speed) were computed using a custom-made software package (see Chen et al. 2015). The temperature was maintained at 25°C, 60% RH and a light intensity of 1.2205 kLux in the flight mill experiments.

### Relationship between JH titer and flight capability of *B. dorsalis*

JHs were assayed from two batches of 15-d-old male and female adults. The first batch of adults was used as a control while the second batch of adults were tethered on the flight mills for 1, 2, 5, 10 and 24 hours following the flight capacity test described in Chen et al. (2015). The JH of each tested insect was determined immediately after the flight test according to the procedures described earlier. Each treatment was replicated six times. 15-d-old male and female adults were used for the study as they were known to have the highest flight ability (see Chen et al., 2015).

### Effects of JH III treatment on the flight capability of *B. dorsalis*

The JH III reference standard was dissolved in acetone (Beijing Chemical Reagent Co., Ltd.) and four different solutions of 0.01 μg/μl, 0.1 μg/μl, 1 μg/μl and 10 μg/μl were prepared. The solutions were stored in a refrigerator at −20°C prior to use. The test insect was anesthetized using crushed ice for 1.5 min. 5 μl of each JH III solution was dripped on the flight muscles of 14-d-old adult male and female using an Eppendorf pipette. Each of the adults in the control group was treated only with 5 μl of acetone. Each treatment was repeated six times. Treatments for the control and JH III-treated groups were applied between 7–8 a.m. The test adults were returned to their rearing cages and tested after 24 h. JH III- and acetone-treated adults were tethered on the flight mills for 13 h to determine their flight duration, distance and average speed (see Chen et al., 2015). The tethered flight tests were conducted under a controlled environment at 25°C, 60% RH and a light intensity of 1.2205 kLux.

### Data analysis

JH titers in adults of different ages, sexes, stages of ovary development, flight hours on the flight mills, flight duration, flight distance and average flight speed were analysed using one-way ANOVA, followed by multiple comparisons using Fisher’s Least Significant Different (LSD) if data are statistically significant. Regression analysis was used to determine the relationship between JH titer and flight duration, flight distance and average flight speed. Data were analysed with SPSS Version 17.0 (SPSS Inc. 2008).

## Results

### JH quantification

The JH III reference standard and JH III extract from the flight muscle of *B. dorsalis* showed stable peaks. The retention times of the JH III standard and JH III from the flight muscle were 10.33 min and 10.39 min, respectively (Fig. 1). The calibration curve y = 38700x − 4.63, R^2^ = 1.00 (x, concentration of standard solution; y, peak area) showed a strong linear relationship between the concentration and peak area of the JH III standard (Fig. 2).

**Figure 1.**
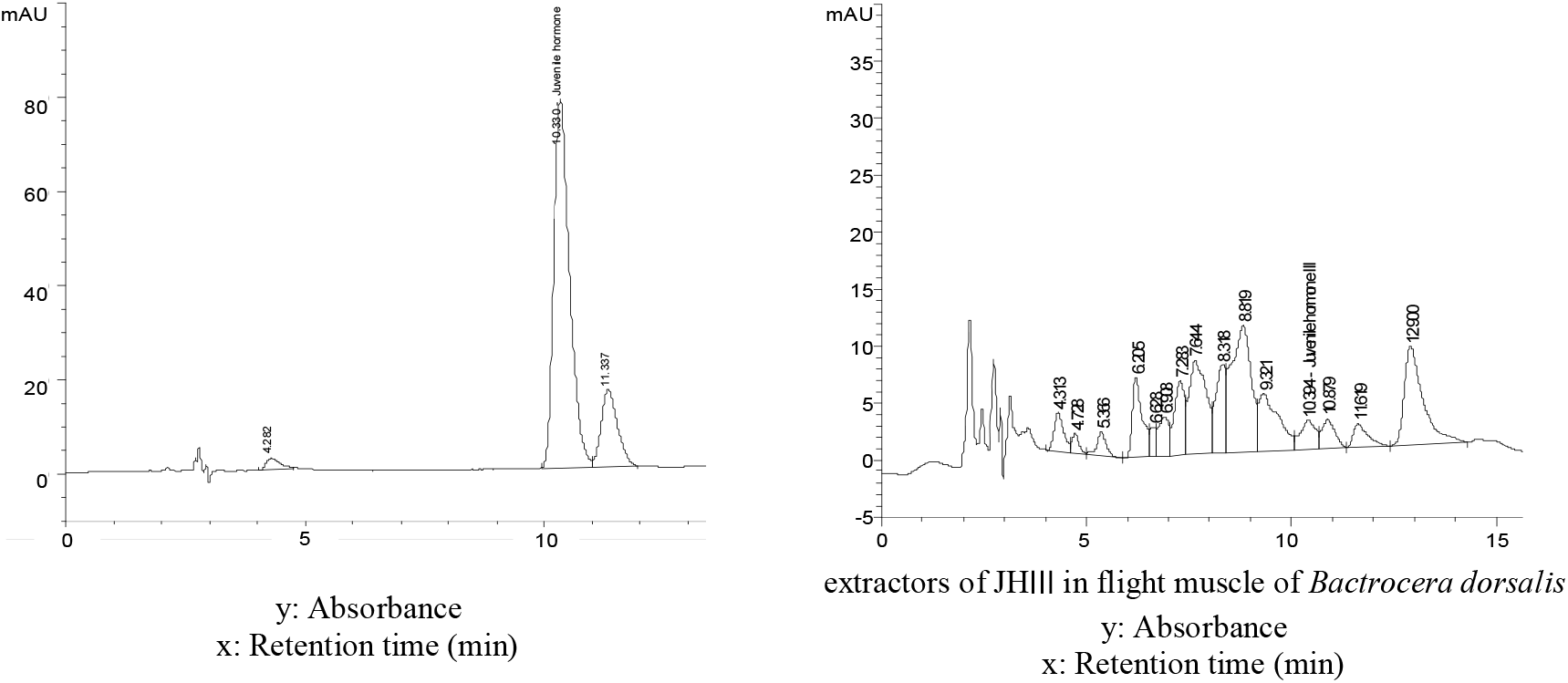
HPLC chromatogram of the juvenile hormone ◻ reference standard and juvenile hormone III extract from the flight muscle of *Bactrocera dorsalis*.

**Figure 2.**
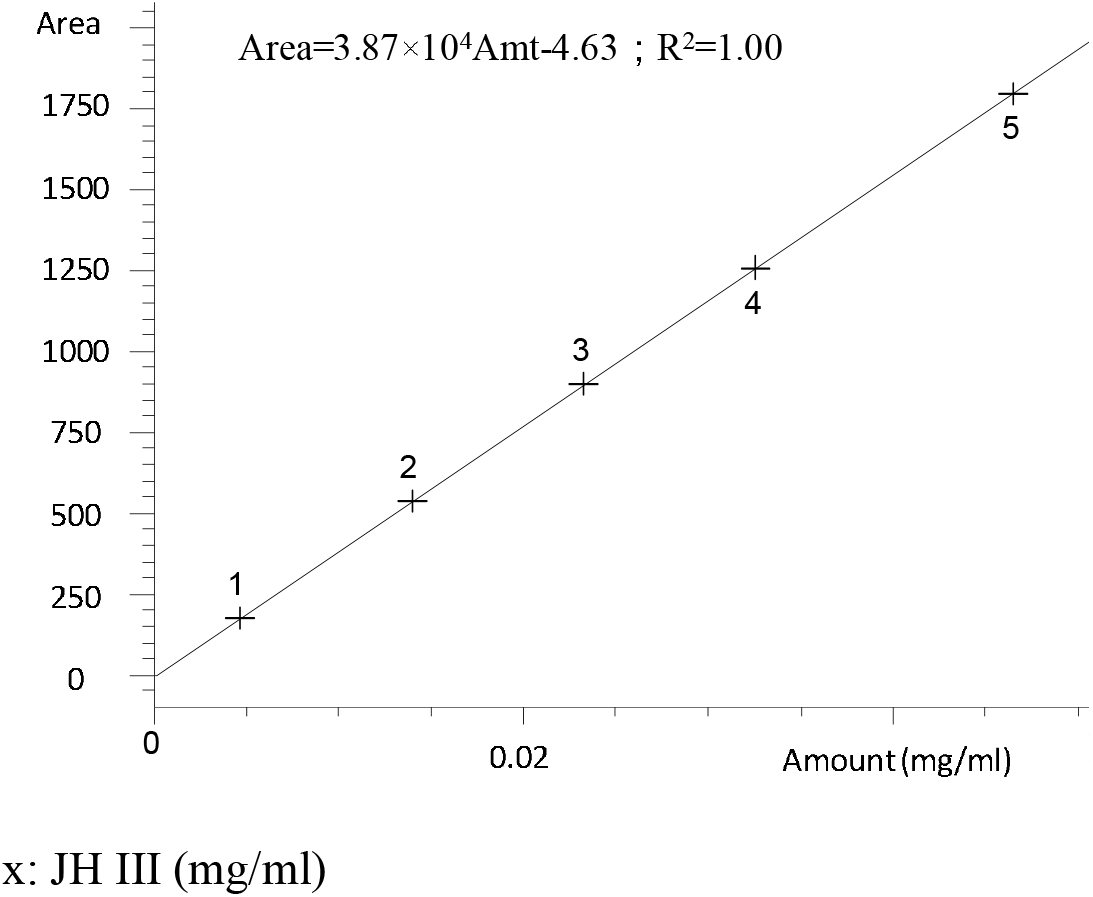
Regression curve of juvenile hormone ◻ reference standard.

### JH titers in *B. dorsalis* adults of different ages and sexes

JH titers in adults of different ages were significantly different (*P* <0.001; *F* = 7.31; *df* = 4). JH titers increased after adult emergences but began to decrease after reaching its peak levels in 15-d-old adults (Fig. 3). The JH titers of 5-, 10-, 15-, 20- and 25-d-old females were 3.15 μg/g, 7.50 μg/g, 8.70 μg/g, 5.73 μg/g and 2.73 μg/g, respectively. The highest JH titer was recorded in 15-d-old female, which was 3.19 times higher than the 25-d-old female. Similarly, after the emergence of male adults, JH titers began to rise and then declined after day 15. The JH titers of 5-, 10-, 15-, 20- and 25-d-old males were 2.68 μg/g, 7.12 μg/g, 8.62 μg/g, 5.56 μg/g and 1.81 μg/g, respectively. JH titer of 15-d-old male was significantly higher than the other age groups and was 4.76 times higher than the 25-d-old male. If comparing the JH titers between the males and females, irrespective of age, overall females have higher JH titers relative to the males of the same age (Fig. 3), but it was not be significant (*P* = 0.0569; *F* = 7.03; *df* = 1).

**Figure 3.**
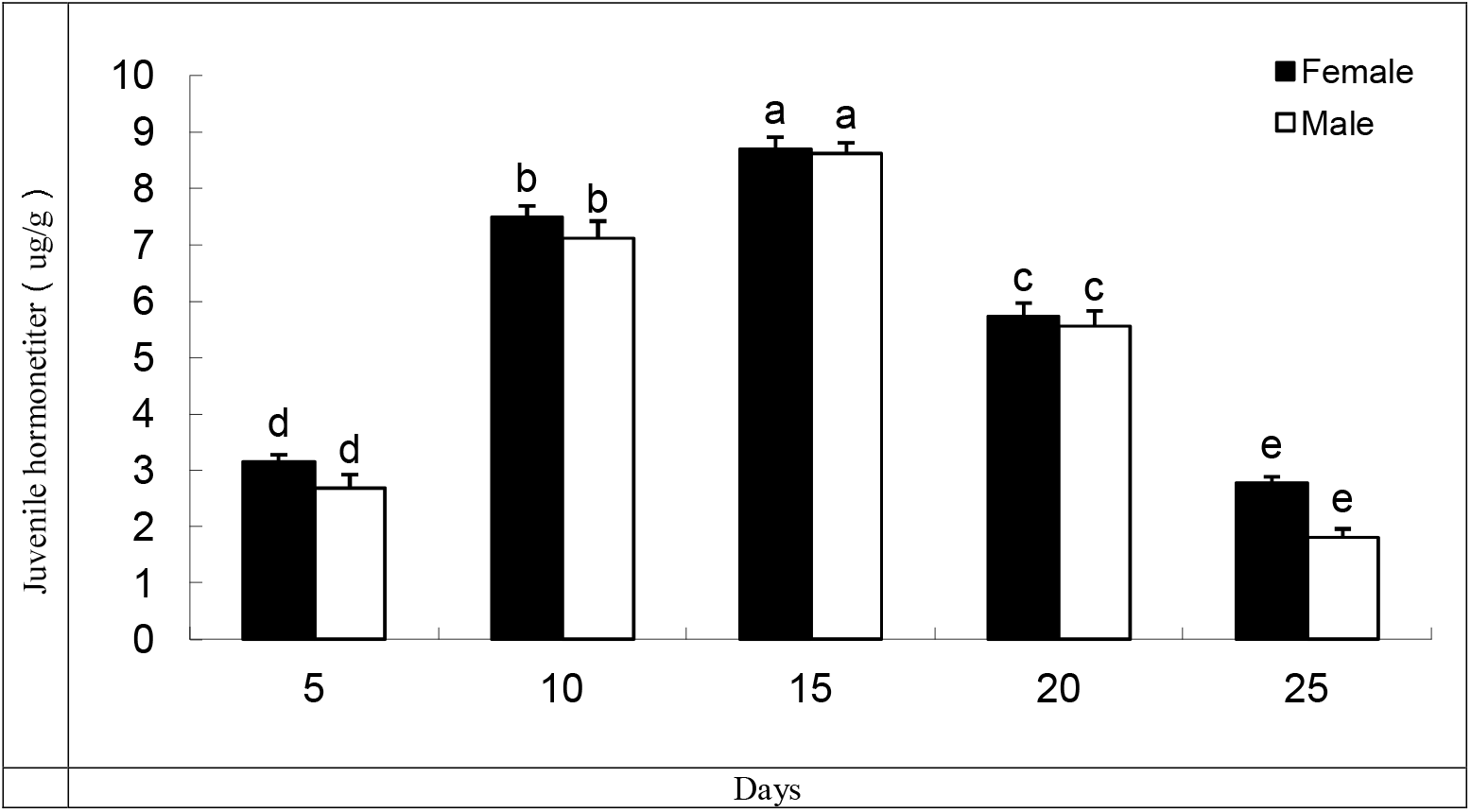
Variations in the juvenile hormone titer in the flight muscle of *Bactrocera dorsalis* adults at different days after adult emergence. Mean ± SE. Bars with different letters represent significant differences at the 5% level. Bars with the same letters are not significantly different at the 5% level (test).

### Relationship between JH titer and flight capability of *B. dorsalis*

The flight duration, distance and average speed of female and male adults increased with the increasing JH titers in the flight muscles of *B. dorsalis* (Fig. 4). JH titer in the flight muscle was positively correlated to the flight duration of females (y = 0.6819 − 0.0401x + 0.0126x^2^, R^2^ = 0.8813) and males (y = 0.6832 − 0.0674x + 0.0147x^2^, R^2^ = 0.7946). Increase in JH titer in the flight muscle had increased the flight distance of females (y = 3.1288 − 1.236x + 0.1302x^2^, R^2^ = 0.8628) and males (y = 1.4993 − 0.5242x + 0.0788x^2^, R^2^ = 0.8794). JH titer also influenced the average flight speed of females (y = 0.5754 − 0.1333x + 0.0179x^2^, R^2^ = 0.7585) and males (y = 0.5481 − 0.1614x + 0.5481x^2^, R^2^ = 0.8497).

**Figure 4.**
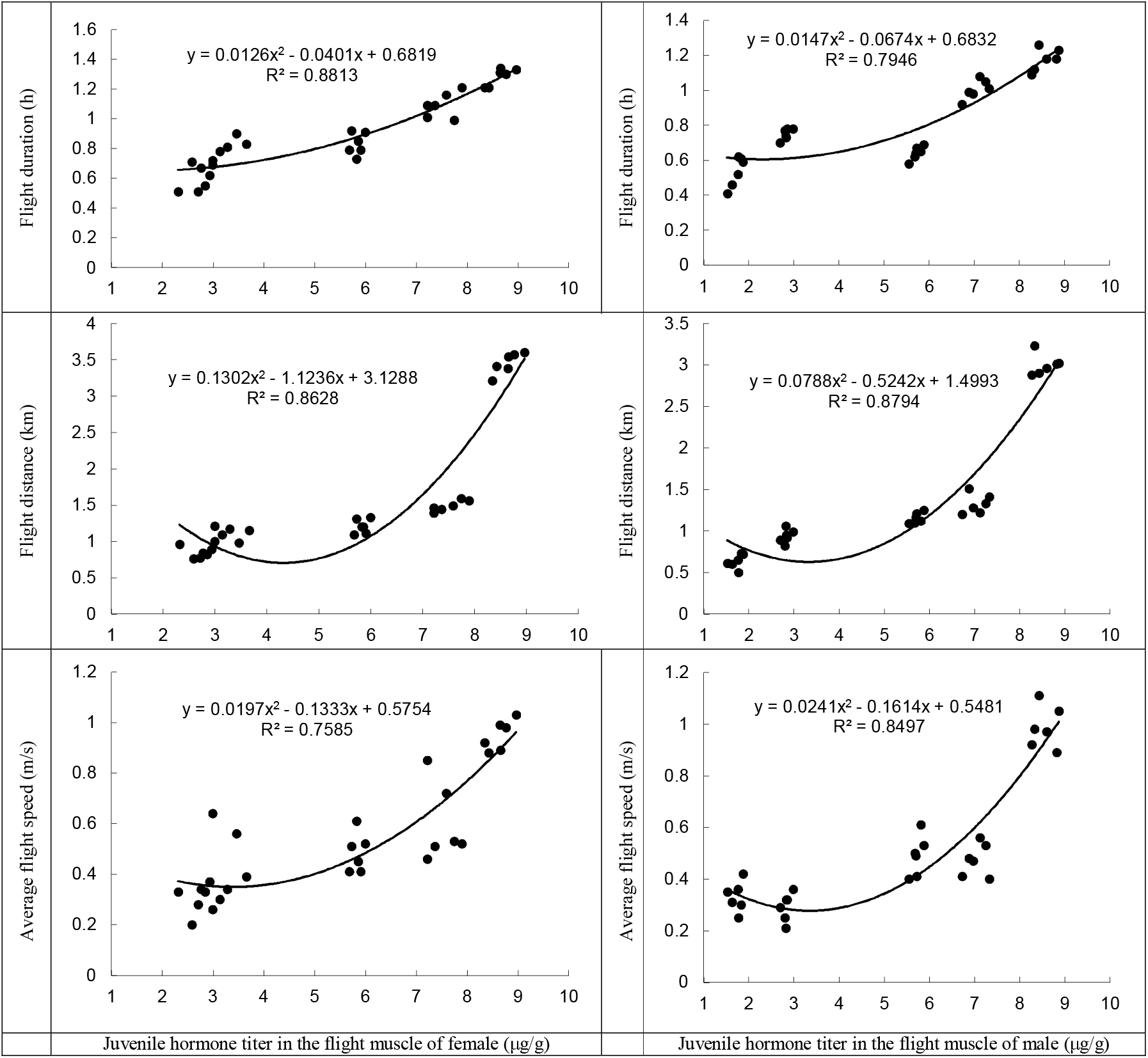
Relationship between juvenile hormone titer in the flight muscle and flight capability of *Bactrocera dorsalis* male and female adults.

### Relationship between JH titer of flight muscle and ovary development in *B. dorsalis*

The results showed that JH titers were significantly different in the different stages of ovary development in *B. dorsalis* (*P* < 0.001; *F* = 1170.51; *df* = 4). JH titer of *B. dorsalis* flight muscle increased and peaked at the expectant stage of mature eggs (stage III) before decreasing (Fig. 5). JH titers of flight muscle at the ovarian developmental stages of I, II, III, IV and V were 3.1406 μg/g, 7.4802 μg/g, 8.7570 μg/g, 5.6702μg/g and 2.7378 μg/g, respectively. JH titer in the stage III of ovarian development was three times higher than stage V (Fig. 5).

**Figure 5.**
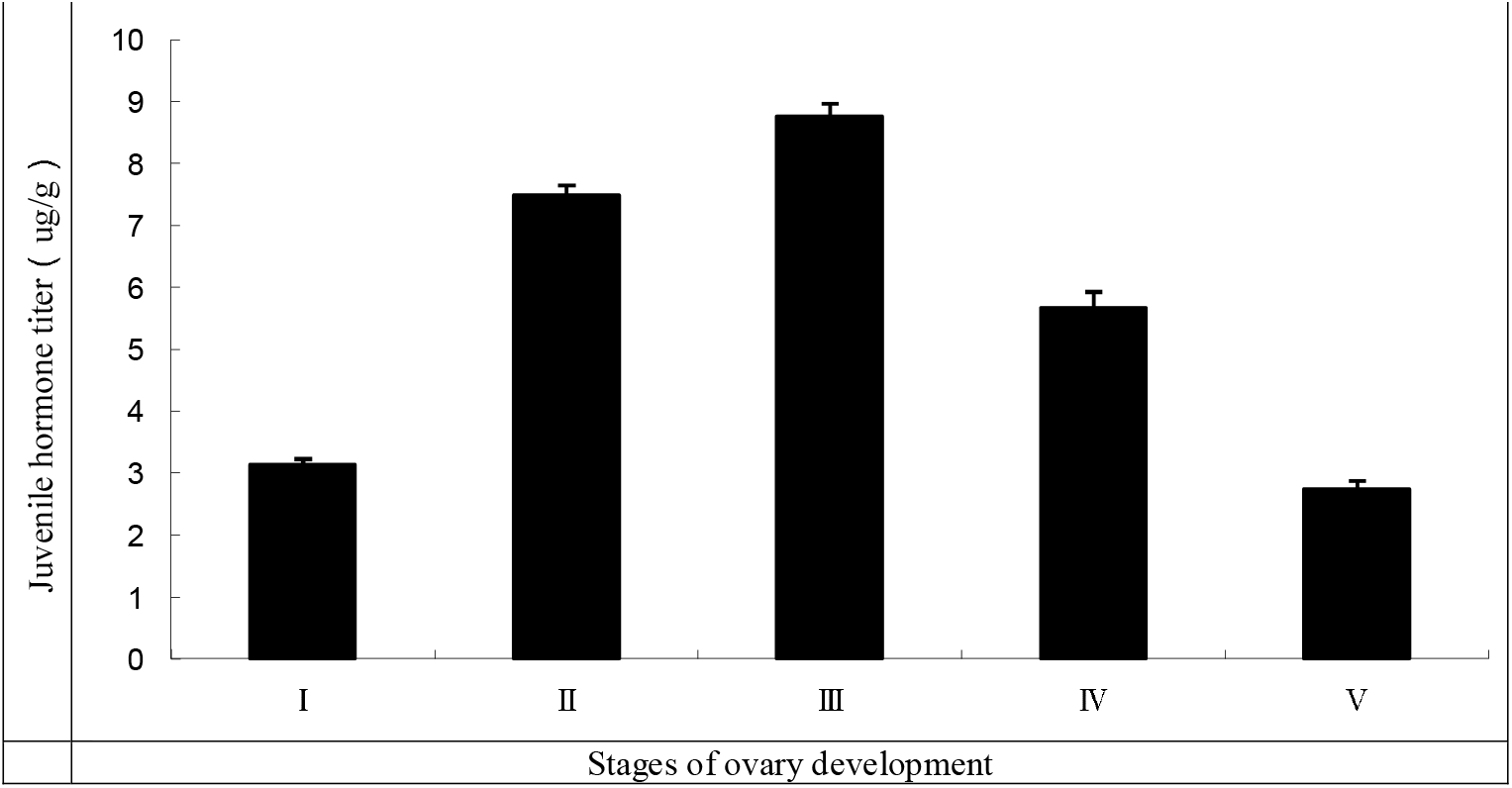
Relationship between the ovarian developmental stages and titers of juvenile hormone ◻ in the flight muscle of *Bactrocera dorsalis* adults. Mean ± SE. Bars with different letters represent significant differences at the 5% level. Bars with the same letters are not significantly different at the 5% level (test).

### Effects of tethered flight on JH titer

Female and male adults had higher JH titers post-flight (*P* < 0.001; *F* = 403.05; *df* = 5) and JH titers of females were significantly higher than the males (*P* < 0.001; *F* = 64.75; *df* = 1). The JH titer of 15-d-old female had increased from 8.62 μg/g to 9.21 μg/g, 9.39 μg/g, 10.07 μg/g, 10.46μg/g, and 11.22 μg/g after 1, 2, 5, 10 and 24 hours of flying, respectively. However, there were no significant differences after 1 and 2 hours of flying although the JH titers had increased 6.38% and 8.00%, respectively. In contrast, there were significant differences in the JH titers of females after 5, 10 and 24 hours on the flight mill, with an increase of 16.00%, 20.41% and 29.93%, respectively (Fig. 6). In 15-d-old male adult, the JH titer had increased from 8.50 μg/g to 9.01 μg/g (4.57% increase), 9.16 μg/g (7.03%), 9.77 μg/g (15.00%), 10.20 μg/g (19.92%), and 11.04 μg/g (29.43%) after 1, 2, 5, 10 and 24 hours of flying, respectively. Likewise, the increases in the JH titers were not significantly different after 1 and 2 hours of flying but were significantly higher with longer duration of tethered flight (Fig. 6).

**Figure 6.**
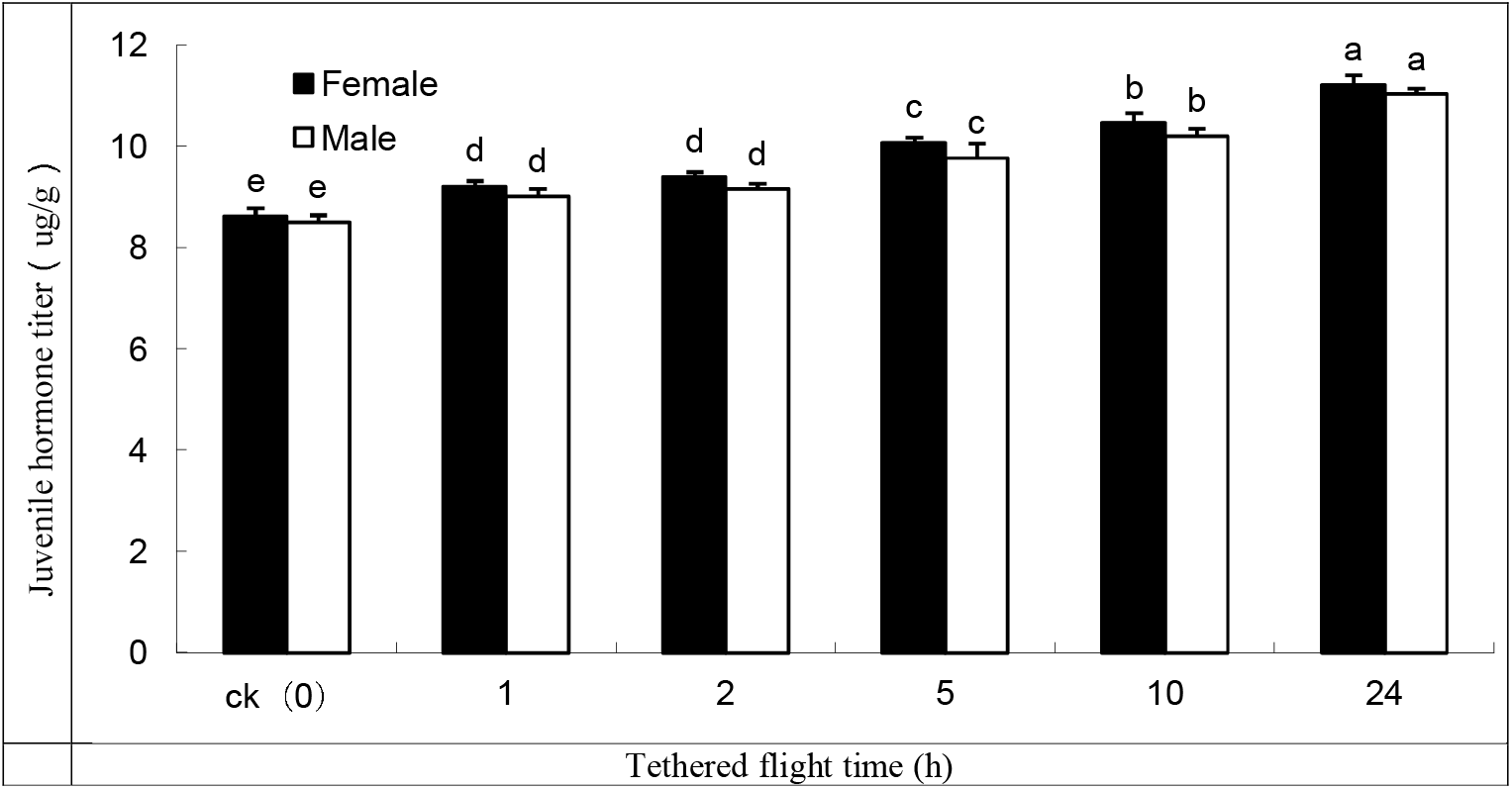
Variations in the juvenile hormone titer in the flight muscle of 15-d-old *Bactrocera dorsalis* adult before and after tethered flight of 1, 2, 5, 10 and 24 hours. Mean ± SE. Bars with different letters represent significant differences at the 5% level. Bars with the same letters are not significantly different at the 5% level (test).

### Effects of tethered flight on flight capability

There were no significant differences in the flight duration (*P* = 0.6906; *F* = 0.16; *df* = 1), flight distance (*P* = 0.1554; *F* = 2.18; *df* = 1) and flight speed (*P* = 0.9059, *F* = 0.01; *df* = 1) between the 15-d-old males and females flown at different hours on the flight mills (Fig. 7) although the females were slightly better fliers than the males. However, the flight duration (*P* < 0.001; *F* = 1020.08; *df* = 4) and distance (*P* < 0.001, *F* = 501.43; *df* = 4) of both male and female adults increased gradually with longer hours spent on the tethered flights. The flight durations of males were 0.16, 0.38, 0.79, 1.08 and 1.41 hours while females recorded 0.15, 0.37, 0.79, 1.08 and 1.47 hours during tethered flights of 1, 2, 5, 10 and 24 hours, respectively (Fig. 7). The flying distance of males were 0.47, 1.10, 2.13, 2.80 and 3.43 km while the females recorded 0.46, 1.09, 2.19, 2.75 and 3.66 km at 1, 2, 5, 10 and 24 hours, respectively (Fig. 7). Both males and females recorded the longest flight duration and distance when flown for 24 hours on the flight mills. However, the average flight speed of male and female adults did not change significantly and was maintained at about 1 m/s in all the tethered flights (*P* = 0.9913; *F* = 0.07; *df* = 4) (Fig. 7).

**Figure 7.**
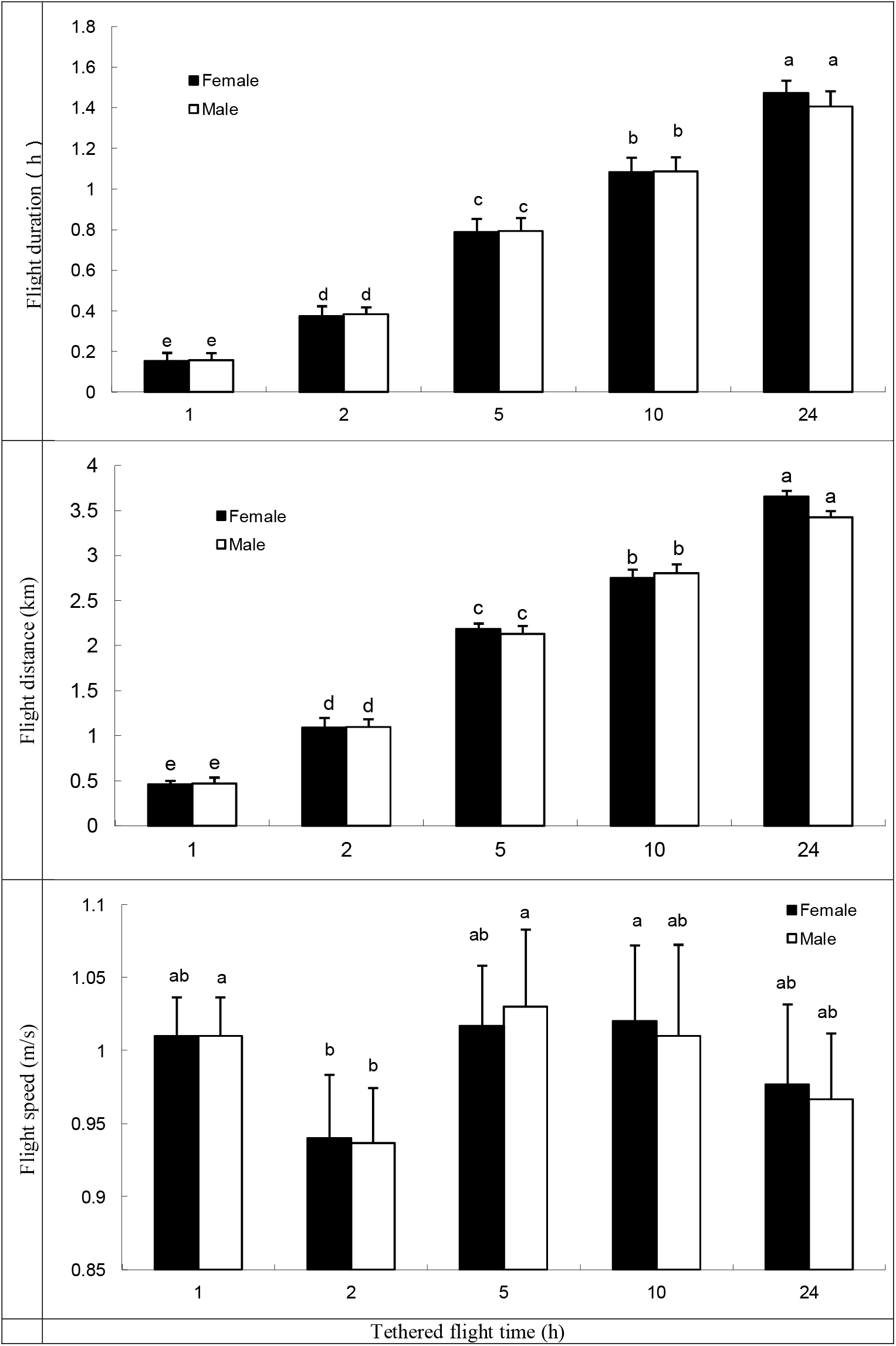
Variations in the flight capability of 15-d-old *Bactrocera dorsalis* female and male adults flown at different flight hours. Mean ± SE. Bars with different letters represent significant differences at the 5% level. Bars with the same letters are not significantly different at the 5% level (test).

### Effects of JH titer on flight capability of *B. dorsalis* on tethered flight

An increase in the JH titer corresponded to the increase in the flight duration and distance for both males and females (Fig. 8). JH titers were positively correlated to flight duration (females, y = −9.4529 + 1.3924x − 0.0372x^2^, R^2^ = 0.9526; males, y = −9.4554 + 1.4494x − 0.0417x^2^, R^2^ = 0.9543) and distance (females, y = −51.612 + 9.1319x − 0.3751x^2^, R^2^ = 0.9378; males, y = −47.159 + 8.5458x − 0.3594x^2^, R^2^ = 0.9543) (Fig. 8). However, the average flight speed was not influenced by JH titer and the regression equation between JH titer and average flight speed could not be established as the model fits the data poorly (Fig. 8).

**Figure 8.**
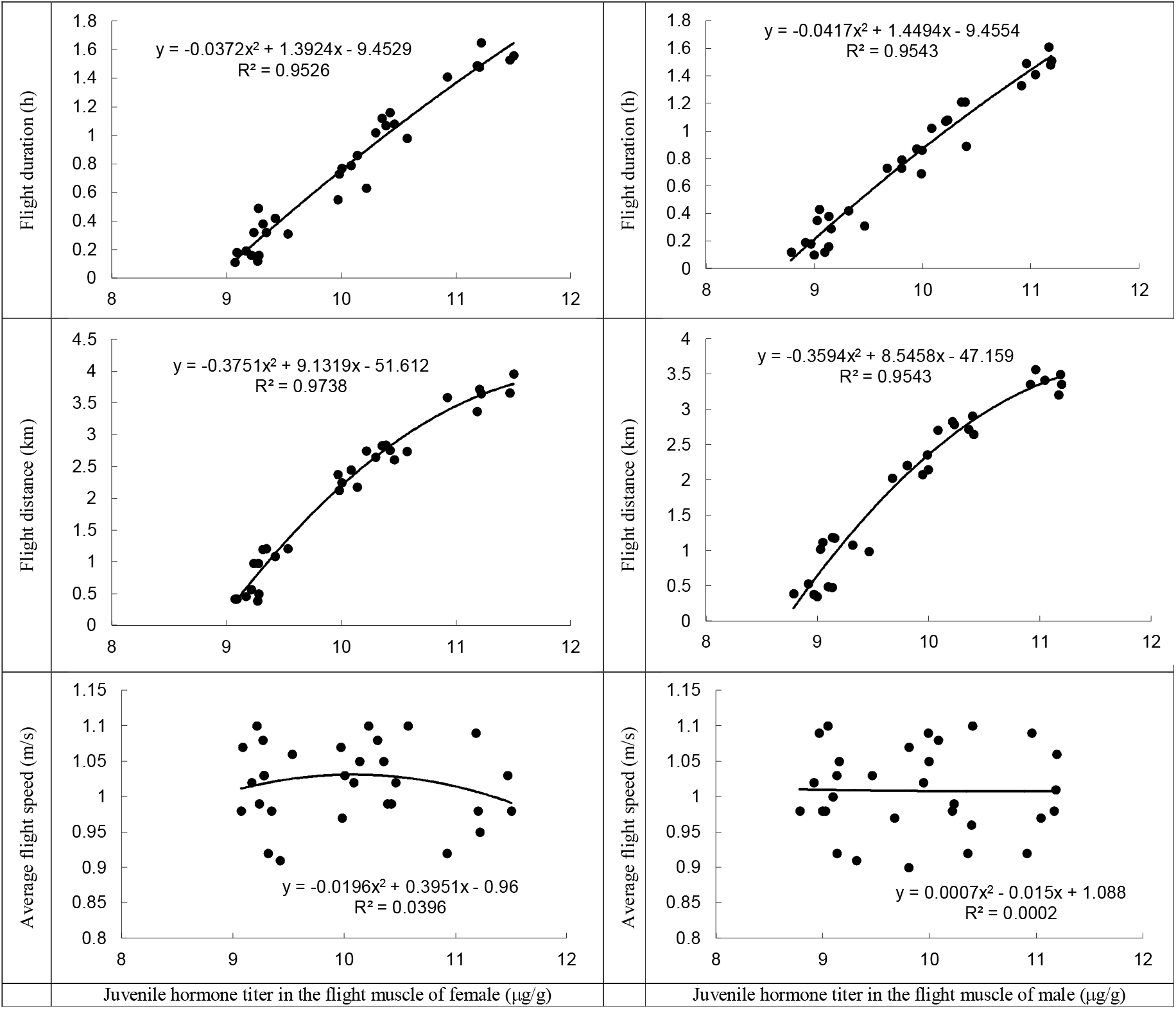
Relationship between flight capability and juvenile hormone titer in the flight muscle of 15-d-old *Bactrocera dorsalis* adult.

### Effects of JH III treatment on the flight muscle of *B. dorsalis*

Flight duration (*P* = 0.0114; *F* = 19.64; *df* = 1) and flight distance (*P* = 0.0049; *F* = 31.68; *df* = 1) improved significantly between the male and female adults treated with JH III compared to adults without treatment. Flight duration (*P* = 0.0087; *F* = 3.82; *df* = 4) and flight distance (*P* = 0.0166; *F* = 12.11; *df* = 4) of both males and females treated in different solutions of JH III differed significantly. However, the average flight speed between JH III-and non-treated female and male adults was not significantly different (*P* = 0.0665; *F* = 6.27; *df* = 1). Likewise, the average flight speed of both female and male adults at different JH III solutions did not differ (*P* = 0.6334; *F* = 0.70; *df* = 4) (Fig. 9).

**Figure 9.**
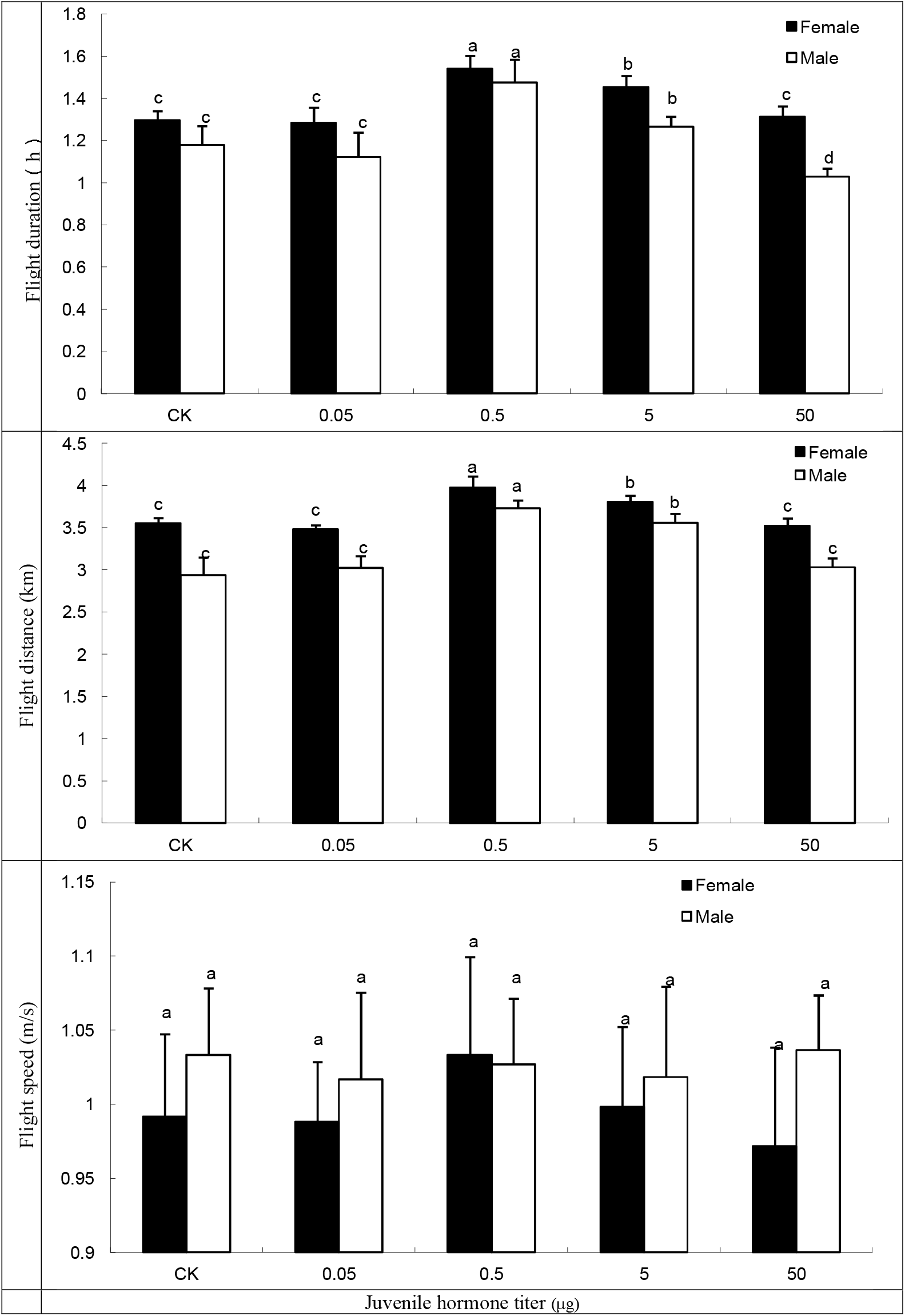
Flight capacity of *Bactrocera dorsalis* treated at different doses of juvenile hormone ◻. Mean ± SE. Bars with different letters represent significant differences at the 5% level. Bars with the same letters are not significantly different at the 5% level (test).

Flight distance and duration of both female and male adults treated with 0.5 μg and 5 μg of JH III increased significantly, compared to those without treatment (Fig. 9). Control females flew for 1.30 hours at the average speed of 0.99 m/s and a distance of 3.55 km, while control males flew for 1.18 hours at the average speed of 1.03 m/s and a distance of 2.94 km. Treatment with 0.5 μg JH III improved the flight capabilities of females (flight duration, 1.54 hours; speed, 1.03 m/s, distance, 3.97 km) and males (flight duration, 1.48 hours; speed, 1.03 m/s; distance, 3.73 km) (Fig.9). There were also significant differences in the flight capabilities of females (flight duration, 1.45 hours; speed, 1.00 m/s; distance, 3.80 km) and males (flight duration, 1.27 hours; speed, 1.02 m/s; distance, 3.56 km) when treatment dose was increased to 5 μg. Meanwhile, adults treated with 0.05 μg and 50 μg of JH III were not significantly different from the control group (Fig.9).

## Discussion

High performance liquid chromatography (HPLC) provides an accurate and efficient bioassay on JHs in insects (Dai et al. 1997a, Dai et al. 1997b, Dai et al. 2001, Ouyang and Li 2003) compare to traditional *Galleria* wax-wound bioassay, radioimmunoassay and chromatography, which have low efficiency and detection rates (Gilbert and Schneiderman 1960, Dahm et al. 1976). The separation conditions in the HPLC vary based on the type of instruments and equipment used (Dai et al. 2001, Wang and Li. 2002, Jiang and Luo 2005). In this study, the ratio of the mobile phase solvent and flow rate were based on the shortest elution time and these separation conditions were comparable to other JH assays such as the assays on the cypress sawfly (Wang and Li. 2002), brown planthopper (Dai et al. 2001) and Oriental armyworm (Jiang and Luo 2005). For the quantification of JH in *B. dorsalis* flight muscle, HPLC separation conditions using methanol: water at the ratio of 80:20 (v/v) and a flow rate of 0.8 ml/min were highly efficient as the retention time of JH III in the flight muscle was 10.39 min relative to the JH III standard of 10.33 min.

The results from this study show that females of *B. dorsalis* contained higher JH titers than males of the same age. The high level of JH corresponded to the hormonal changes required for ovarian development and maturation in female adults (Fu and Chen 1984, Flatt et al. 2005, Harshman and Zera 2007, Chen 2013). At the initiation of ovarian development in the Pacific beetle cockroach *Diploptera punctata* (Eschscholtz), the CA is stimulated to secrete more JH for the synthesis and secretion of yolk protein in the oocytes. The rise and fall of the JH titer after ovary maturation were also observed in the large milkweed bug (Rankin and Riddiford 1978). Therefore, this explains the differences in hormonal requirements and concentrations between the male and female of *B. dorsalis*.

JH titer varies with the developmental stages of the insect (Wang 2001; Rauschenbach et al. 2007, Goodman and Granger 2005). The high JH titers in the final instar larva of the cypress sawfly *Chinolyda flagellicornis* (F. Smith) and brown planthopper *Nilaparvata lugens* (Stål) were believed to maintain the larval characteristics during the larval growth (Dai et al. 2001, Wang and Li 2002). Larval diapause in the orange wheat blossom midge *Sitodiplosis mosellana* (Géhin) was influenced by JH and the overwintering larva had the highest JH titer than the other stages of development (Li et al. 2006). The change of JH titer with insect growth and development also occurred in the East Asian migratory locust whereby the JH titer increased continuously after eclosion and peaked when the adult reached 10-d of age, which also coincided with its peak ovarian development, before gradually decreasing subsequently (Liu 2007).

JH promotes development of flight muscles but inhibits flight activity at high concentrations to compensate for an insect’s maturation processes, such as ovarian development (Rankin and Riddiford 1978). High JH titer was associated with high flight capability (duration and distance) and longer flying hours stimulated the production and higher levels of JH III in *B. dorsalis*. In the study on the long-winged sand cricket, Zhang et al. (2011) detected higher JH titer in the flight muscle post-flight. The emigrant and immigrant populations of the Oriental armyworm *Mythimna separata* (Walker) also exhibited differing levels of JH. The immigrating population with higher levels of JH was attributed to the long flight duration, which stimulated the production of JH (Jiang and Luo 2005).

In this study, *B. dorsalis* adults treated with moderate concentration 0.5 μg to 5 μg of JH III had improved flight activities with longer flight durations and distance covered but when the dose was too low or too high outside the range of significant doses, no effect was observed. In migratory insects, JH treatment was observed to enhance the flight behavior and ovarian development of the convergent lady beetle *Hippodamia convergens* Guérin-Méneville (Rankin and Rankin 1980) while JH suppression led to the impairment of both migration and reproduction abilities of the shield bug *Eurygaster integriceps* Puton (Polivanova and Triseleva 1985).

JH titer in the flight muscle of *B. dorsalis* was closely related to its age and sex. A well-developed 15-d-old *B. dorsalis* adult contained the highest level of JH while the lowest JH level was detected in the initial stage of eclosion (~ 5-d-old) and towards the end of the adult life span (~ 25-d-old) (Chen et al. 2015). Based on our previous study, laboratory-reared *B. dorsalis* adults usually lived for 25–30 d. As reported by Chen et al. (2015), 15-d-old adult female has the strongest flight capacity due to its well-developed flight muscle structure, and based on the findings from this study, we can conclude that JH correlates to the development of flight muscle after adult emergence. The female also tends to have a higher JH titer in its flight muscle to synchronize with the development of its reproductive system. Similarly, JH titer peaked during the stage III of ovarian development, but was the lowest in the early and towards the end of its ovarian development.

Flight activity stimulates the secretion of JH and causes an increase in the JH titer in the flight muscle. On the other hand, the increase in the JH titer corresponds to the increase in the flight duration and distance of *B. dorsalis*, suggesting that JH and flight activity have a mutually reinforcing relationship. These findings improved our understanding on the relationship between JH in flight muscle and flight ability, including the physiological roles of JH on the flight activities of *B. dorsalis*. Metabolic and behavioral activities of an insect is regulated by a variety of hormones involving complex mechanisms, therefore further experimental studies are needed to elucidate the internal mechanism and processes of different regulating hormones on the flight activities of *B. dorsalis* in future.

## Acknowledgments

This research was supported by the National Natural Science Foundation Program of P.R. China (31660208, 31160162), Joint Special Agriculture Foundation Program of Yunnan (2017FG001-024), the Central Financial Forestry Science and Technology Promotion Demonstration Project (YUN[2017]TG08) and Research Team Construction Project of Yunnan Academy of Forestry (Research Team of Forest Diseases and Pests Control).

